# Sulfate adenylyl transferase kinetics and mechanisms of metabolic inhibitors of microbial sulfate respiration

**DOI:** 10.1101/2021.03.29.436835

**Authors:** Hans K. Carlson, Matthew D. Youngblut, Steven A. Redford, Adam J. Williamson, John D. Coates

## Abstract

Sulfate analog oxyanions that function as selective metabolic inhibitors of dissimilatory sulfate reducing microorganisms (SRM) are widely used in ecological studies and industrial applications. As such, it is important to understand the mode of action and mechanisms of tolerance or adaptation to these compounds. Different oxyanions vary widely in their inhibitory potency and mechanism of inhibition, but current evidence suggests that the sulfate adenylyl transferase/ATP sulfurylase (Sat) enzyme is an important target. We heterologously expressed and purified the Sat from the model SRM, *Desulfovibrio alaskensis* G20. With this enzyme we determined the turnover kinetics (k_cat_, K_M_) for alternative substrates (molybdate, selenate, arsenate, monofluorophosphate, and chromate) and inhibition constants (K_I_) for competitive inhibitors (perchlorate, chlorate, and nitrate). These measurements enable the first quantitative comparisons of these compounds as substrates or inhibitors of a purified Sat from a respiratory sulfate reducer. We compare predicted half-maximal inhibitory concentrations (IC_50_) based on Sat kinetics with measured IC_50_ values against *D. alaskensis* G20 growth and discuss our results in light of known mechanisms of sensitivity or resistance to oxyanions. This analysis helps with the interpretation of recent adaptive laboratory evolution studies and illustrates the value of interpreting gene-microbe-environment interactions through the lens of enzyme kinetics.

## Main text

Selective inhibitors of dissimilatory sulfate reducing microorganisms (SRM) are both valuable tools for ecological studies and treatment strategies to abrogate unwanted sulfide production in industrial systems ^1,2^. The best studied selective SRM inhibitors are the inorganic oxyanion sulfate analogs including molybdate, tungstate, selenate, chromate, monofluorophosphate, arsenate, nitrate, perchlorate, and chlorate ^3–6^. Remarkably, although direct interaction with the sulfate activating enzyme, sulfate adenylyl transferase (Sat) has been implicated as a primary mode of action for these compounds ^3,7^, there is little kinetic data for these sulfate analogs as substrates or inhibitors of purified Sat enzymes from respiratory SRM. Other mechanisms of toxicity, tolerance and adaptation have been implicated for oxyanions against model SRM including competition for sulfate uptake ^8,9^, ATP consuming futile cycles ^10,11^and detoxification via enzymatic reduction and efflux ^5,12^.

We heterologously expressed, purified and kinetically characterized the Sat from the model SRM, *Desulfovibrio alaskensis* G20 as reported previously ^13^. Alongside our previous measurements with molybdate ^13^, we report kinetic parameters for four other alternative substrates including selenate, arsenate, monofluorophosphate, and chromate ^13–15^ (Table 1A). Previously we found that perchlorate is a competitive inhibitor of the Sat ^13^, and we now report inhibition constants (K_I_) for the other competitive inhibitors nitrate and chlorate, both of which are competitive with the oxyanion substrate, molybdate, used in our assays but not competitive with the Sat co-substrate, ATP (Table 1B).

**Table 1.** Kinetic parameters for *D. alaskensis* G20 Sulfate adenylyl transferase (Sat)

A.
| Substrate | $k_{cat}$ ($s^{-1}$) | $K_M$ (mM) | $k_{cat}/K_M$ ($M^{-1} s^{-1}$) | Reference |
| --- | --- | --- | --- | --- |
| $MoO_4^{2-}$ | $14.5 \pm 1.2$ | $3.26 \pm 0.55$ | $4.4 \times 10^3$ | <sup>13</sup> |
| $SeO_4^{2-}$ | $0.284 \pm 0.032$ | $1.77 \pm 0.55$ | $1.6 \times 10^2$ | This work |
| $AsO_4^{2-}$ | $3.88 \pm 0.32$ | $0.0617 \pm 0.026$ | $6.3 \times 10^4$ | This work |
| $FPO_4^{2-}$ | $11.0 \pm 1.6$ | $1.44 \pm 0.56$ | $7.6 \times 10^3$ | This work |
| $CrO_4^{2-}$ | $16.0 \pm 0.79$ | $0.793 \pm 0.14$ | $2.0 \times 10^4$ | This work |

B.
| Inhibitor | $K_i$ - Varied $MoO_4^{2-}$ (mM) | $K_i$ - Varied ATP (mM) | Average $K_i$ (mM) | Reference |
| --- | --- | --- | --- | --- |
| $ClO_4^-$ | $0.138 \pm .014$ | $0.915 \pm 0.15$ | 0.5265 | <sup>13</sup> |
| $ClO_3^-$ | $0.316 \pm 0.017$ | $1.45 \pm 0.076$ | 0.833 | This work |
| $NO_3^-$ | $3.60 \pm 0.27$ | $12.5 \pm 0.98$ | 8.05 | This work |

While this set of sulfate analogs has previously been inferred to target Sat based on their selective influence on SRM metabolic activity and growth ^4,5,7,10^, cell lysate enzyme assays ^3^, genetic screens ^7,16^ and induction of the sulfate reduction regulon ^7^ our results with the purified enzyme enable the first quantitative ranking of these compounds as substrates or inhibitors of Sat. It is striking that the relative affinities (K_M_ and K_I_) of these oxyanions vary by over an order of magnitude given their similar geometries and formal charges (Table 1). The highest affinity substrate is arsenate (K_M_= 0.0617 ± 0.026) while the lowest affinity substrate is molybdate (K_M_ = 3.26 ± 0.55). The highest affinity inhibitor is perchlorate (K_I_ *=* 0.138 ± 0.014) while the lowest affinity inhibitor is nitrate (K_I_ *=* 3.60 ± 0.27).

The potency of a competitive substrate or inhibitor against cellular growth is determined by the affinity of the inhibitor for its primary target and detoxification reactions. For cytoplasmic enzymes, mechanisms of transport and efflux are important to consider. Thus, we can compare the measured affinities against Sat with known inhibitory potencies of these oxyanions against growth of *D. alaskensis* G20 ^5^ to infer the extent to which Sat is a primary target of these oxyanions and the magnitude of other processes that influence intracellular inhibitor concentration or other modes of toxicity. While inhibitor K_I_ or K_M_ is independent of the substrate concentration, enzyme IC_50_ increases at higher substrate concentrations for competitive substrates and inhibitors (Equation 1) ^17^.

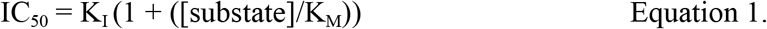

Thus, from measurements of sulfate K_M_ and competitive inhibitor K_I_ or competitive substrate K_M_, we can estimate the competitive substrate/inhibitor IC_50_ against Sat for a given concentration of sulfate (Figure 1). To model IC_50_s we use a sulfate K_M_ of 2.93 ± 0.26 mM empirically determined for a homologous *Desulfovibrio* Sat ^18^. The homologous Sat is 81% identical to the G20 Sat, and molybdate K_M_ values only vary between 1 mM and 3 mM for different *Desulfovibrio* ^*19*^. Thus, this is likely a reasonable approximation for the G20 Sat K_M_. Because we do not know the cytoplasmic concentrations of sulfate during batch growth in the presence of these inhibitors we calculated Sat IC_50_s at three concentrations of sulfate including: 15 mM sulfate (the mean extracellular concentration in G20 batch cultures ^5,9,12,13,20^), 5 mM sulfate (the mean intracellular sulfate concentration in typical G20 batch cultures ^21^), and 0.15 mM sulfate (which is close to the lowest intracellular concentration measured in G20 batch cultures ^21^ and an order of magnitude lower than the Sat sulfate K_M_).

**Figure 1.**
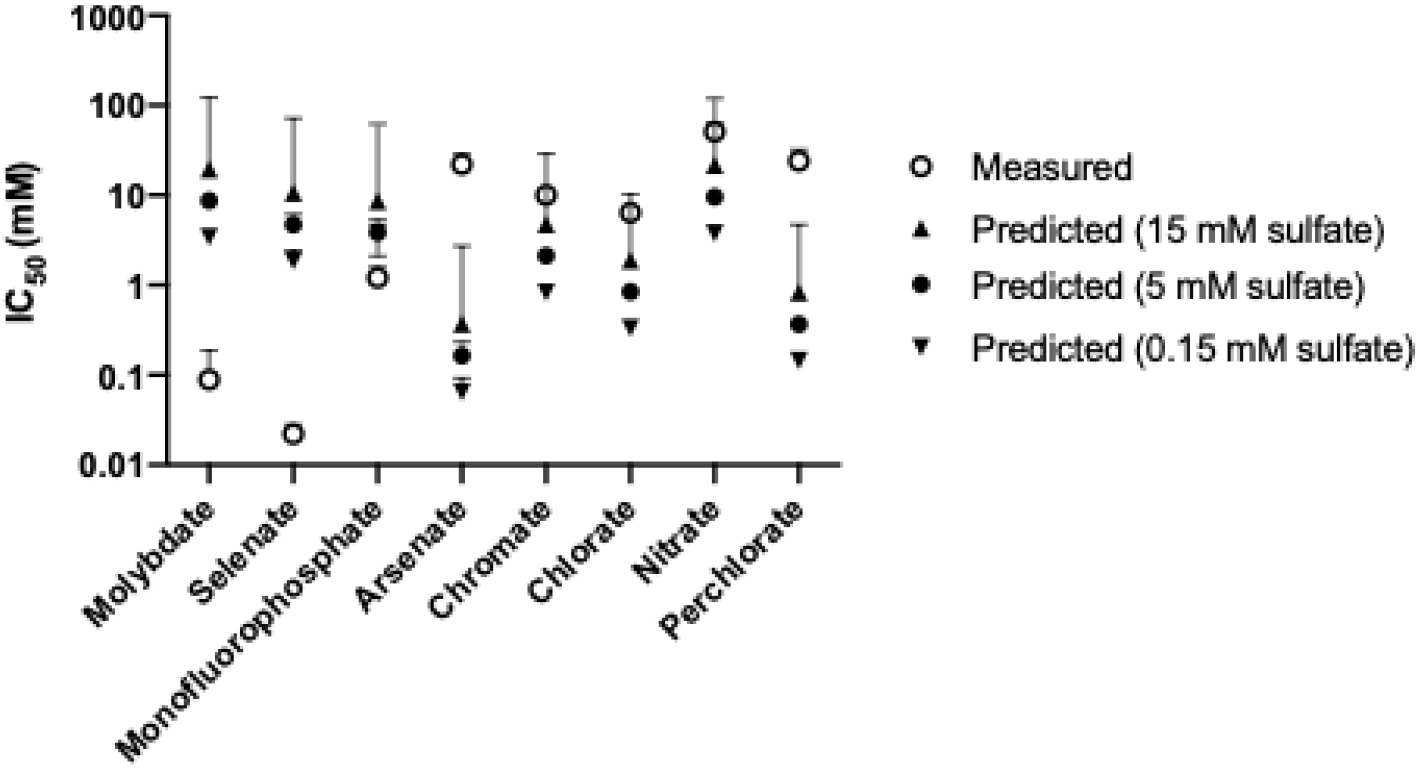
Measured and Predicted IC_50_ values for *D. alaskensis* G20 Sat. Measured IC_50_s are from Reference 5 and error bars represent the 95% confidence intervals of dose—response fits. Predicted IC_50_s are based on measured kinetic parameters for for the G20 sulfate adenyltransferase (Table 1) using Equation 1 and calculated using for intracellular sulfate concentrations of 15 mM (closed upright triangles) 5 mM (closed circles), and 0.15 mM (closed inverted triangles). Predicted IC_50_s error bars are upper and lower bounds calculated based on error in Sat kinetic measurements range (this work, references 13,18).

For the Sat substrates molybdate, selenate and monofluorophosphate, the predicted Sat IC_50_s are higher than the measured growth IC_50_s. As such, the measured IC_50_s must reflect an additional mode of growth inhibition apart from activity as competitive substrates of Sat or bioconcentration via uptake with minimal efflux of the inhibitor in the G20 cytoplasm. Molybdate and selenate form unstable APS analogs as products of Sat catalyzed reactions which rapidly decompose ^22,23^. This drives a non-productive, “futile” cycle which leads to rapid ATP hydrolysis and is thought to be a major mode of cellular toxicity of these compounds ^10^. Selenate and molybdate are also competitive with sulfate uptake ^8^ and, at least in *D. vulgaris* Hildenborough, an additional protein aside from Sat may catalyze non-productive adenosine 5’-phosphomolybdate formation and ATP hydrolysis, but it is unknown if similar proteins are in *D. alaskensis* G20 ^11^. Monofluorophosphate reacts with ATP at the Sat as a dead-end substrate to form the stable product ADPβF ^24^, so ATP consumption through futile cycling is not a likely mechanism of cellular inhibition. However, fluoride ion (F^-^) toxicity may contribute to the monofluorophosphate mechanism of action, but this is ameliorated by a fluoride efflux pump ^5^.

The other divalent oxyanions in our panel, arsenate and chromate, are less inhibitory to *D. alaskensis* G20 growth than predicted based on the Sat IC_50_. Arsenate is the highest affinity Sat substrate from our panel with a Sat K_M_ 50 to 100-fold lower than molybdate or selenate (Table 1, Figure 1A). The measured arsenate IC_50_ against G20 growth is ∼10-fold higher than the predicted arsenate Sat IC_50_ and ∼100-fold higher than the measured molybdate or selenate IC_50_s. However, arsenate is known to be catalytically reduced and effluxed by G20 and deletion of the arsenate reductase and efflux systems renders G20 ∼10-fold more sensitive to arsenate ^12^. Arsenate also reacts abiotically with sulfide ^25^. These observations are consistent with the difference the measured arsenate IC_50_ being lower than predicted (Figure 1). Chromate is the second highest affinity Sat substrate in our panel, and the predicted IC_50_ is slightly lower than the measured IC_50_. Chromate is catalytically reduced by G20 ^26^ and reacts abiotically with sulfide ^27^ which will increase the effective concentration required to inhibit Sat. Taken together, our results are consistent with Sat being an important target of both arsenate and chromate in G20, and this is consistent with the previous observation that both of these compounds are selective inhibitors of SRM in marine enrichment cultures ^5^.

We also compared predicted Sat IC_50_s with measured growth IC_50_s for the competitive inhibitors perchlorate, chlorate, and nitrate (Figure 1). The mechanism of action of these compounds against SRM in complex natural systems is primarily due to bio-competitive exclusion via growth of nitrate or perchlorate respirers and the production of reactive nitrogen and potentially chlorine species ^1,28,29^. However, at higher concentrations, these oxyanions are direct competitive inhibitors of the sulfate reduction pathway ^7^. Understanding the targets and adaptation mechanisms to competitive inhibitors may aid in the development of next-generation small molecule inhibitors that are selective against SRM ^30^.

For nitrate and chlorate, predicted IC_50_s are less than 10-fold lower than the measured IC_50_s while the predicted perchlorate IC_50_ is nearly 50-fold lower than the measured IC_50_. Apart from competitive inhibition of Sat, cellular permeability, reactivity and efflux can influence the measured growth IC_50_s, cytoplasmic reactivity will make cells more sensitive to these compounds through the generation of reactive nitrogen and chlorine species (RNS/RCS). While perchlorate is kinetically very stable, chlorate and nitrate generate cytoplasmic RNS/RCS in G20 ^7,16^. No chlorate or nitrate efflux mechanisms in G20 are known and reduction or these compounds is minimal ^7^. Thus, for these compounds it is most likely that measured growth IC_50_s are higher than are expected based on Sat kinetics because oxyanion transport keeps cytoplasmic concentrations of these compounds low.

Indeed, adaptive laboratory evolution and genetics indicate that changes in the permeability of cells to sulfate, nitrate and perchlorate influence sensitivity to these oxyanions. Mutants that increase the expression of sulfate transporters with low perchlorate affinity are implicated in perchlorate resistance ^9^, and loss of function mutants in putative thiosulfate transporters with high nitrate affinity are implicated in nitrate resistance ^7,16,20^. Notably, point mutants that alter the activity of sulfate transporters were not observed suggesting that these compounds target a cytoplasmic enzyme, such as Sat, rather than sulfate uptake. In some perchlorate adapted cultures, point mutants in the Sat emerge that alter the K_i_, clearly indicating that Sat is under selection ^9,13^.

Further characterization of transport kinetics, rates of reduction and efflux, and cytoplasmic concentrations will enable more quantitative predictions of the inhibitory potency and genetic targets of oxyanion inhibitors of sulfate respiration. Nevertheless, our results are consistent with Sat as an important target of these compounds. More generally, comparing the affinities of metabolic inhibitors for transport enzymes versus cytoplasmic metabolic enzymes will be essential for understanding the evolutionary landscapes that microorganisms navigate across environmental gradients of toxicants or nutrients ^31–33^. The extent to which cell surface versus cytoplasmic enzymes are under selection depends on enzyme affinities and the relative concentrations of substrates and inhibitors. Thus, enzyme kinetics measurements are critical for understanding genotype-phenotype relationships in complex environments where a bacterial cell is confronted with physicochemically similar substrates and inhibitors such as metal ions, carbon sources or vitamins.

